# *Plasmodium falciparum* diacylglycerol acyltransferase maintains phospholipid homeostasis to regulate sexual differentiation, ER stress, and cytoadhesion

**DOI:** 10.1101/2025.10.10.681613

**Authors:** Minako Yoshida, Junpei Fukumoto, Shinya Miyazaki, Megumi Tanaka, Suzumi M. Tokuoka, Hideru Obinata, Takaya Sakura, Yoko Onizuka, Daniel Ken Inaoka, Kiyoshi Kita, Kazuhide Yahata, Eizo Takashima, Hideo Shindou, Fuyuki Tokumasu

## Abstract

*Plasmodium falciparum* is the causative agent of human malaria, a life-threating infectious disease that imposes a major global health burden. Lipid metabolism is indispensable for this parasite’s replication and survival, yet most of the molecular components and mechanisms involved remain poorly understood. In eukaryotes, lipid droplets (LDs) serve as dynamic organelles that store neutral lipids (NLs), buffer lipotoxic stress, and regulate signaling pathways, with their biogenesis controlled by diacylglycerol o-acyltransferases (DGATs). Although *P. falciparum* encodes a putative DGAT (PF3D7_0322300), its role in the parasite life cycle has not been elucidated. We generated conditional PfDGAT-knockout parasites to investigate the enzyme’s functional significance. PfDGAT deficiency led to parasite death, accompanied by reduced LD formation, elevated phospholipid levels, and induction of ER stress. Moreover, PfDGAT deletion altered protein trafficking, resulting in the decreased cytoadherence of parasite-infected erythrocytes to human brain microvascular endothelial cells, and suppressed parasite sexual differentiation. Thus, PfDGAT deletion affected multiple aspects of the parasite’s life cycle, highlighting its critical role in parasite survival and pathogenesis. Our findings provide new insights into parasite lipid homeostasis and highlight DGAT as a potential target of antimalarial intervention.

## Introduction

Malaria parasites invade erythrocytes and undergo exponential proliferation during intraerythrocytic developmental cycles, producing dozens of daughter merozoites within a single round. This proliferative process greatly expands the parasite membrane surface area, necessitating a large supply of phospholipid species. Although the parasites can incorporate certain lipids from the host and external environment, the availability of de novo-synthesized and stored phospholipids is essential for their ability to adapt to rapid fluctuations in lipid content while maintaining an appropriate lipid composition^1–3^. The sexual differentiation of the parasite into the gametocyte stage also requires a characteristic lipid profile. Therefore, lipid synthesis is important for their survival in the bloodstream, subsequent uptake by mosquitos, and adaptation to the new environment within the mosquito^4–6^.

The relative composition of lipid species directly determines both the biophysical properties and biological functions of cellular membranes, including the maintenance of membrane integrity, the formation of vesicles and their fusion to a target membrane, and the functional activities of membrane proteins. Lipid species with various acyl chain lengths and head groups have distinct molecular shapes and exhibit different intermolecular interactions, and thereby generate membrane lipid heterogeneities characteristic to specific cell types and organelles. In vesicular protein trafficking, the budding and fusion of small vesicles involve the formation of an intermediate structure (“hemifusion”) characterized by negative membrane curvature^7,8^. To generate and maintain such curvature, phospholipids with small head groups, such as phosphatidylserine (PS), phosphatidylethanolamine (PE), phosphatidic acid, and the more rectangular diacylglycerol (DAG), must be present at defined concentrations to enable efficient hemifusion^9,10^. A comprehensive understanding of the enzymes that synthesize these lipid classes, which are critical determinants of vesicle formation, is crucial, as proper protein trafficking cannot otherwise be explained.

Triacylglycerols (TAGs), a major group of NLs, constitute the core of lipid droplets (LDs), together with cholesteryl esters^11,12^. Although LDs primarily function as reservoirs of NLs, they also protect against lipotoxicity by ameliorating ER stress, serve as principal sites of metabolite flux, act as nutrient sensors through LD-localizing proteins, and form membrane contact sites (MCSs) with other organelles^13^. These functions are essential for maintaining cellular homeostasis^14^. In eukaryotes, TAGs are synthesized in two ways: acyl chain incorporation into DAG from the *de novo* phospholipid synthesis (Kennedy) pathway, catalyzed by diacylglycerol acyltransferase (DGAT), and (acyl-CoA-independent) acyl chain transfer from phospholipids to DAG by phospholipid:diacylglycerol acyltransferase^15^. Mammals possess two DGAT enzymes, DGAT1 and DGAT2, which share no sequence homology but compensate for each other functionally. Both enzymes are localized to the ER, while DGAT2 is also localized to LDs^16^. DGAT1 has relatively broad substrate specificity, enabling it to act on several substrates^17^, whereas DGAT2 functions primarily as the canonical DGAT enzyme catalyzing TAG synthesis. DGAT2-knockout mice exhibit impaired skin barrier function, leading to postnatal death, and display a decrease in carcass TAG levels of up to 90%^18^. By contrast, DGAT1-knockout mice survive, maintain a lean body mass, and are resistant to high-fat-diet-induced obesity^19^. In murine embryonic fibroblasts, DGAT1 deficiency, which impaired TAG synthesis, promotes oleate-induced lipotoxicity^20^.

*P. falciparum* database homology searches have identified a single putative DGAT gene, designated *Pfdgat* (PF3D7_0322300). The PfDGAT protein shares sequence similarity with human DGAT1 at the amino acid level, but not with DGAT2. It has been predicted to be essential based on a double-crossover recombination study and piggyBac genome-wide screening^21,22^. In addition, ghosts of *P. falciparum*-infected erythrocytes, as well as the heterologous expression of PfDGAT in CHO cells, demonstrated diacylglycerol acyltransferase activity, suggesting that PfDGAT may function as a DGAT enzyme in malaria parasites^21,23^. However, the detailed role of PfDGAT in parasite biology remains unexplored.

We therefore generated a conditional PfDGAT-knockout parasite and demonstrated that this enzyme is essential for asexual proliferation. Loss of PfDGAT led to elevated phospholipid levels, which likely suppressed parasite sexual differentiation. In addition, PfDGAT dysfunction triggered ER stress, impaired protein trafficking, and prevented parasite cytoadherence to brain microvascular endothelial cells. The broad impacts of this single-gene knockout highlight the importance of the coordinated maintenance of both glycerol backbones and fatty acid chains in *P. falciparum* biology.

## Results

### PfDGAT possesses conserved DGAT1 motifs and the membrane bound *O*-acyltransferase motif

The amino acid sequence of PfDGAT was shown to share 24%–30% similarity with DGAT1s from mammals, flies, nematodes, and plants, but no significant similarity to the DGAT2s from these organisms^21^. To assess the spatial similarity between PfDGAT and human DGAT1 (HsDGAT1), we superimposed the predicted three-dimensional conformations (Fig. 1a). Multiple structural regions overlapped between the two proteins, including the FYXDWWN and the SXXXHEY motifs, both of which are highly conserved among DGAT1 enzymes and essential for catalytic function^24,25,26^. By contrast, the N-terminal intrinsically disordered region (IDR) of HsDGAT1 (residues 1–94) had no corresponding region in PfDGAT, which instead possessed its own distinct IDR between residues 158 and 220 (Fig. 1a). The FYXDWWN and SXXXHEY motifs are conserved in DGATs from *Plasmodium* species but absent in those from the other apicomplexan parasites *Toxoplasma gondii* and *Cryptosporidium parvum* (Fig. 1b). This suggests that these organisms either possess unidentified DGAT(s) or their DGATs feature unique catalytic motifs.

**Fig. 1.**
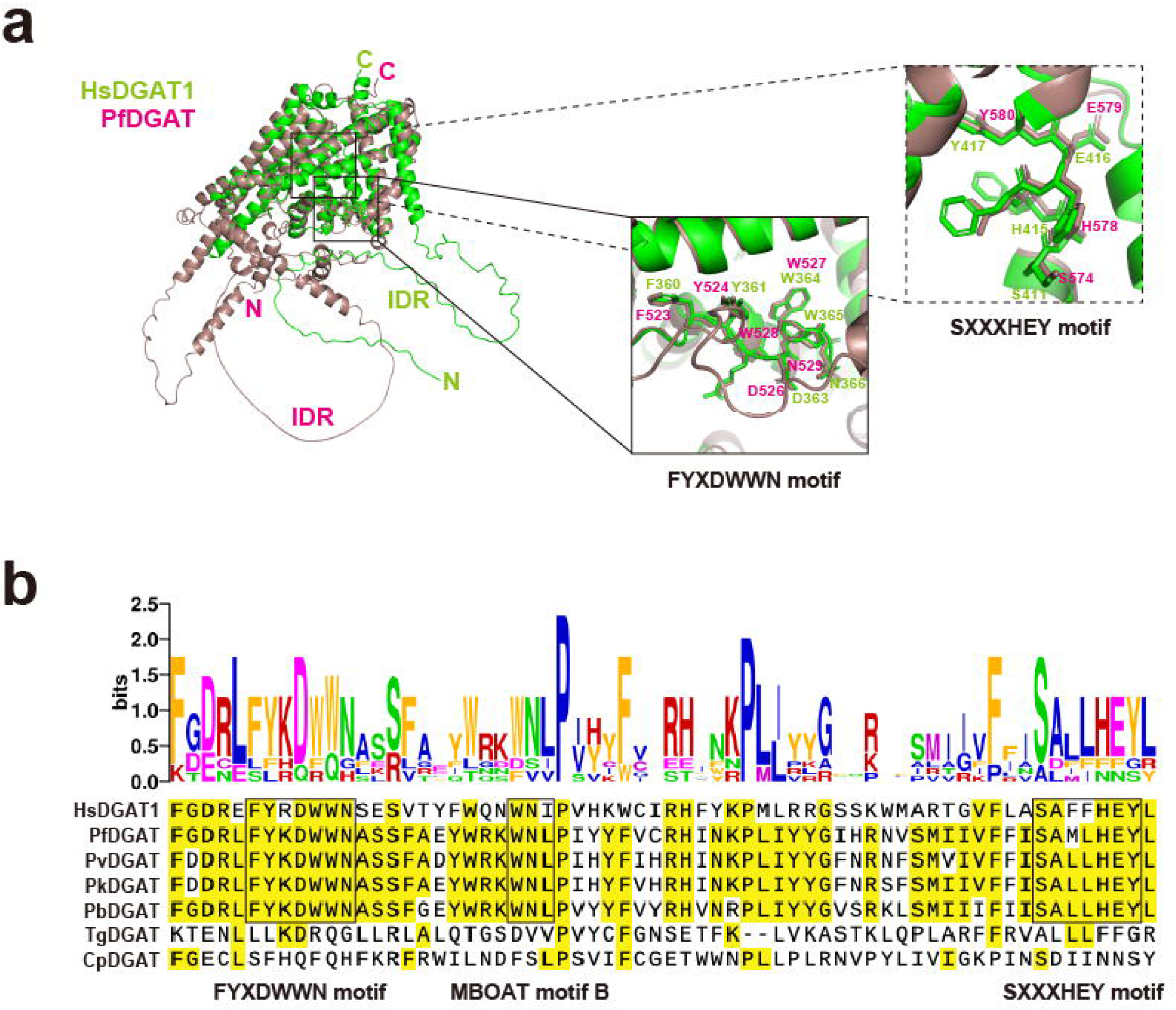
PfDGAT contains conserved DAGT1 functional motifs and the characteristic MBOAT motif. (a) Superimposed three-dimensional structures of PfDGAT and HsDGAT1. Structural data were obtained from the AlphaFold Protein Structure Database. The FYXDWWN and SXXXHEY motifs are shown as the sticks, and the other regions as the cartoon; the motifs are magnified in the right insets. (b) Multiple sequence alignment of DGAT1 orthologs from humans and apicomplexan parasites. Amino acid residues shared in ≥ 50% of the analyzed organisms are highlighted in yellow. The FYXDWWN and SXXXHEY motifs, as well as the MBOAT motif B, are enclosed in black rectangles. Acidic, basic, polar, nonpolar, aromatic, and sulfur-containing amino acids are shown as magenta, red, green, blue, orange, and purple letters, respectively, in the upper sequence logo.

DGAT1 is a member of the membrane-bound *O*-acyltransferase (MBOAT) family characterized by distinct sequence motifs^27,28^. In mammals, DGAT1 contains the MBOAT motif B, characterized by the conserved amino acid sequence “WNI”, whereas the other MBOAT motifs (A, C, and D) are not present^27^. PfDGAT was found to possess WNL at the corresponding position (Fig. 1b), suggesting that PfDGAT belongs to the MBOAT family, similar to mammalian DGAT1.

### PfDGAT is essential for asexual replication and stage transition in *P. falciparum*

To investigate the physiological roles of PfDGAT, which is predicted to function in the Kennedy pathway (Fig. 2a), we generated conditional knockout parasite lines (*Pfdagt*:LoxPint:HA) with the dimerizable Cre recombinase (DiCre) system (Fig. 2b)^29,30^. We constructed a plasmid to conditionally remove the *Pfdgat* sequences, including regions encoding the FYXDWWN and SXXXHEY motifs, in a rapamycin-dependent manner by flanking them with LoxP sites. The plasmid was incorporated into the parasite genome utilizing the selection-linked integration (SLI) method,^31^ and its incorporation was verified by diagnostic PCR (Fig. 2c). To confirm whether the conditional knockout system would function in the established parasite lines, PCR and western blotting were performed (Fig. 2d,e). DNA band shifts were observed in rapamycin-treated *Pfdagt*:LoxPint:HA B10 and G12 clonal lines, which indicated that the *Pfdagt* locus flanked by LoxP sites was excised from the genome (Fig. 2d). Western blot analysis revealed a substantial decrease in PfDGAT expression levels after 24 hours of rapamycin treatment in both clonal lines (Fig. 2e). The loss of PfDGAT in the two synchronized clonal lines resulted in significantly slower growth from day 6 after rapamycin treatment (Fig. 2f). This demonstrated that *Pfdgat* is essential for parasite growth in the erythrocytic stages, consistent with previous reports^21,22^. The *Pfdgat*:LoxPint:HA B10 clonal line was used for further analyses, as no obvious differences were observed between B10 and G12 clones in terms of gene knockout efficiency or growth kinetics.

**Fig. 2.**
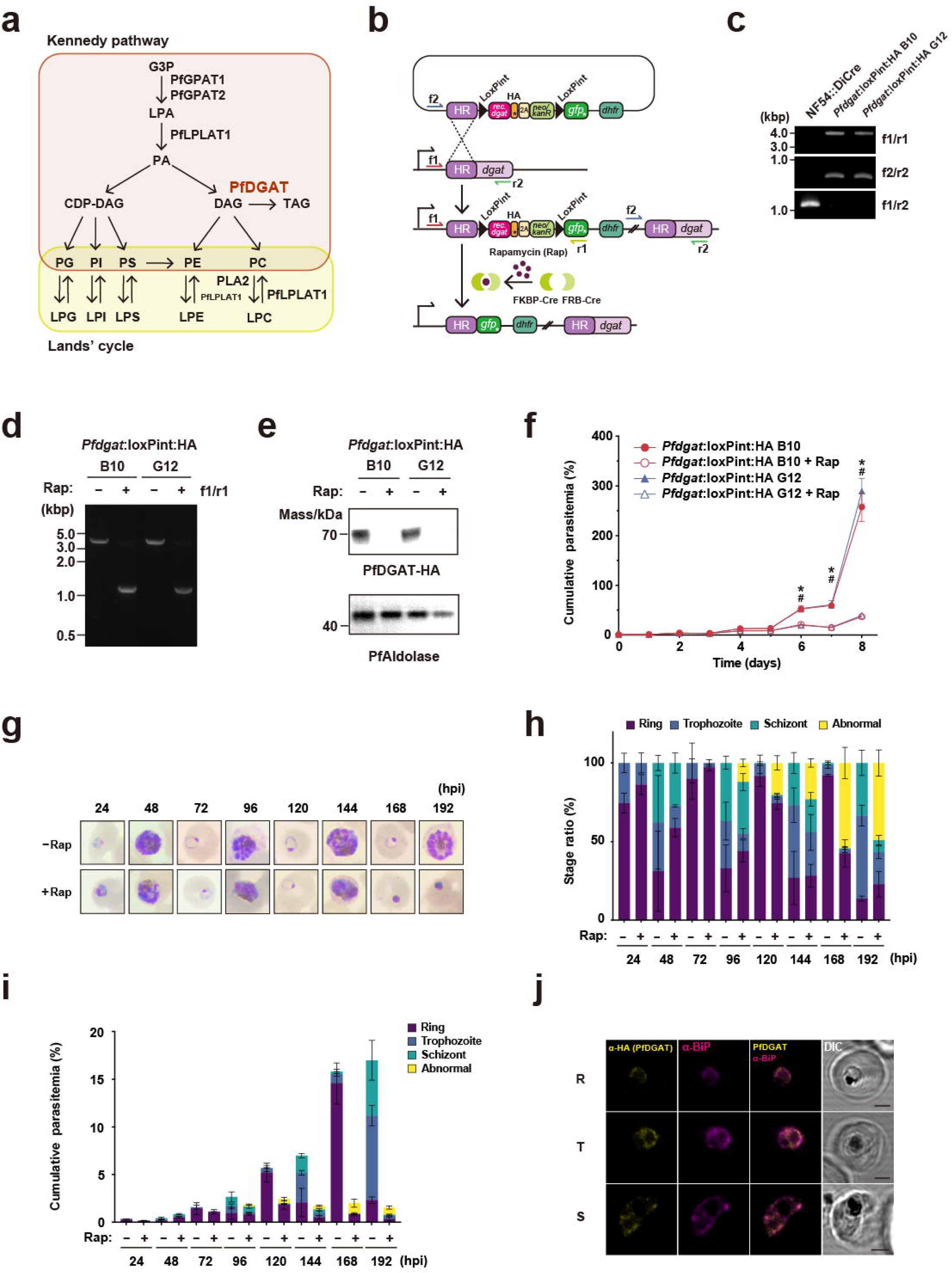
Establishment of PfDGAT conditional knockout *P. falciparum* lines. (a) Putative phospholipid metabolic pathways in *P. falciparum.* PfLPLAT1, *P. falciparum* lysophospholipid acyltransferase 1; the first LPLAT identified in this parasite, functions in both the Kennedy pathway and the Lands’ cycle^61^. G3P, glyceraldehyde 3-phosphate; LPA, lysophosphatidic acid; PA, phosphatidic acid; CDP-DAG, cytidine diphosphate diacylglycerol; PG, phosphatidylglycerol; PI, phosphatidylinositol; PS, phosphatidylserine; PE, phosphatidylethanolamine; PC, phosphatidylcholine; LPG, lysophosphatidylglycerol; LPI, lysophosphatidylinositol; LPS, lysophosphatidylserine; LPE, lysophosphatidylethanolamine; LPC, lysophosphatidylcholine; PfGPAT1 and PfGPAT2, *P. falciparum* glycerol-3-phosphate acyltransferase 1 and 2; PfLPLAT1, *P. falciparum* lysophosphatidic acid acyltransferase 1; and PLA2, phospholipase A2. (b) Design of *Pfdgat* conditional gene knockout with the DiCre system. The 5’ homologous region of *Pfdgat* contains 654 base pairs, beginning from the start codon. Upon rapamycin treatment, the recodonized region, flanked by LoxP introns and containing the FYXDWWN and SXXXHEY motifs, was excised from the parasite genome. Arrows indicate the transcription initiation site and the primer binding sites for PCR-based tests. (c) Diagnostic PCR showed that the target gene was successfully replaced with the targeting constructs: DNA segments were correctly amplified with f1/r1 (NF54::DiCre, not amplified; *Pfdagt*:LoxPint:HA B10 and G12 clones, 3704 bp), f2/r2 (NF54::DiCre, not amplified; *Pfdagt*:LoxPint:HA B10 and G12, 756 bp), and f1/r2 (NF54::DiCre, 1077 bp; *Pfdagt*:LoxPint:HA B10 and G12, not amplified) primer sets. (d, e) Conditional gene knockout of *Pfdagt* was confirmed by PCR and western blotting. (d) The size of DNA amplicons for the 5’ segment shifted from 3704 bp to 1089 bp in *Pfdagt*:LoxPint:HA B10 and G12 after 24 hours of rapamycin treatment. (e) Decreased expression of PfDAGT was detected in *Pfdagt*:LoxPint:HA B10 and G12 with anti-HA antibody after 24 hours of rapamycin treatment. PfAldolase was used as an internal protein control. Theoretical molecular masses (kDa) of HA-tagged PfDAGT and PfAldolase are 81.6 and 40.0, respectively. Experiments were repeated three times with similar results. (f) Growth curve of +Rap or −Rap *Pfdagt*:LoxPint:HA B10 and G12. Data are shown as mean ± SD from n = 3 independent biological replicates. P values were calculated using two-way ANOVA followed by Tukey’s HSD test. \**P* < 0.0001 (−Rap vs. +Rap *Pfdagt*:LoxPint:HA B10); ^#^*P* < 0.0001 (−Rap vs. +Rap *Pfdagt*:LoxPint:HA G12). Experiments were repeated twice with similar results. (g–i) Stage distributions of +Rap or −Rap *Pfdagt*:LoxPint:HA B10. (g) Representative images of Giemsa-stained blood films at described time points. (h, i) Bar charts showing the parasite stage distributions (h) and parasitemia of each stage (i) at the described time points (24, 48, 72, 96, 120, 144, 168, and 192 hours of post-infection [hpi]). Data are shown as mean ± SD from n = 3 independent biological replicates. (j) Representative confocal microscope images of *Pfdagt*:LoxPint:HA B10 co-stained for PfDAGT and ER-resident protein BiP. Samples were immunostained for PfDAGT using an anti-HA antibody (yellow) and for ER with an anti-BiP antibody (magenta). Scale bars = 2 μm. Source data are provided as a Source Data file.

To investigate the stage-dependent effect of PfDGAT deficiency, we tightly synchronized parasites within a 0–4 hour window and monitored their growth over time following rapamycin treatment (Fig. 2g–i). At 96 hours post-infection (hpi), slightly condensed and pyknotic parasites were observed in the rapamycin-treated (+Rap) group, and their proportion had increased to approximately 50% by 168 hpi. No obvious shifts in parasite stages were observed, even at 192 hpi, suggesting that *Pfdgat* deletion had negative effects on all parasite stages or no specific stage-dependent effects.

In mammalian cells and yeast, DGAT family proteins localize at the ER membrane. Ectopically expressed PfDGAT was shown to colocalize with ER-resident chaperone proteins calnexin on the ER^1^, but it is not clear if PfDGAT naturally localizes at the ER. This encouraged us to investigate the subcellular localization of PfDGAT in *P. falciparum*. Immunofluorescence assay (IFA) showed that PfDGAT colocalized with the ER-resident chaperone protein BiP on the ER of the parasite throughout the asexual stage (Fig. 2j), indicating that PfDGAT functions at the ER, as seen for the other DGAT family proteins, in processes such as LD biogenesis.

### PfDGAT deficiency led to a reduction in NL level and induced ER stress in the parasite

Previous reports have described that erythrocytes parasitized with *P. falciparum* and mammalian cells expressing humanized PfDGAT exhibit DGAT activity, synthesizing TAGs from DAGs^1,21^. To determine whether PfDGAT is functionally involved in TAG synthesis within the parasite, we stained NL-rich compartments, such as LDs, with LipidTOX and monitored NL levels in PfDGAT-deficient parasites at 96, 144, and 192 hpi. As sterol esters are absent in malaria parasites^32^, the LipidTOX signal primarily reflects the TAG content. At 96 and 144 hpi, +Rap and rapamycin-untreated (−Rap) parasites showed comparable levels of NL; however, +Rap parasites exhibited a significant reduction in NL levels at 192 hpi (Fig. 3a, b). This suggests that PfDGAT is capable of synthesizing TAG within the parasite during its asexual stage.

**Fig. 3.**
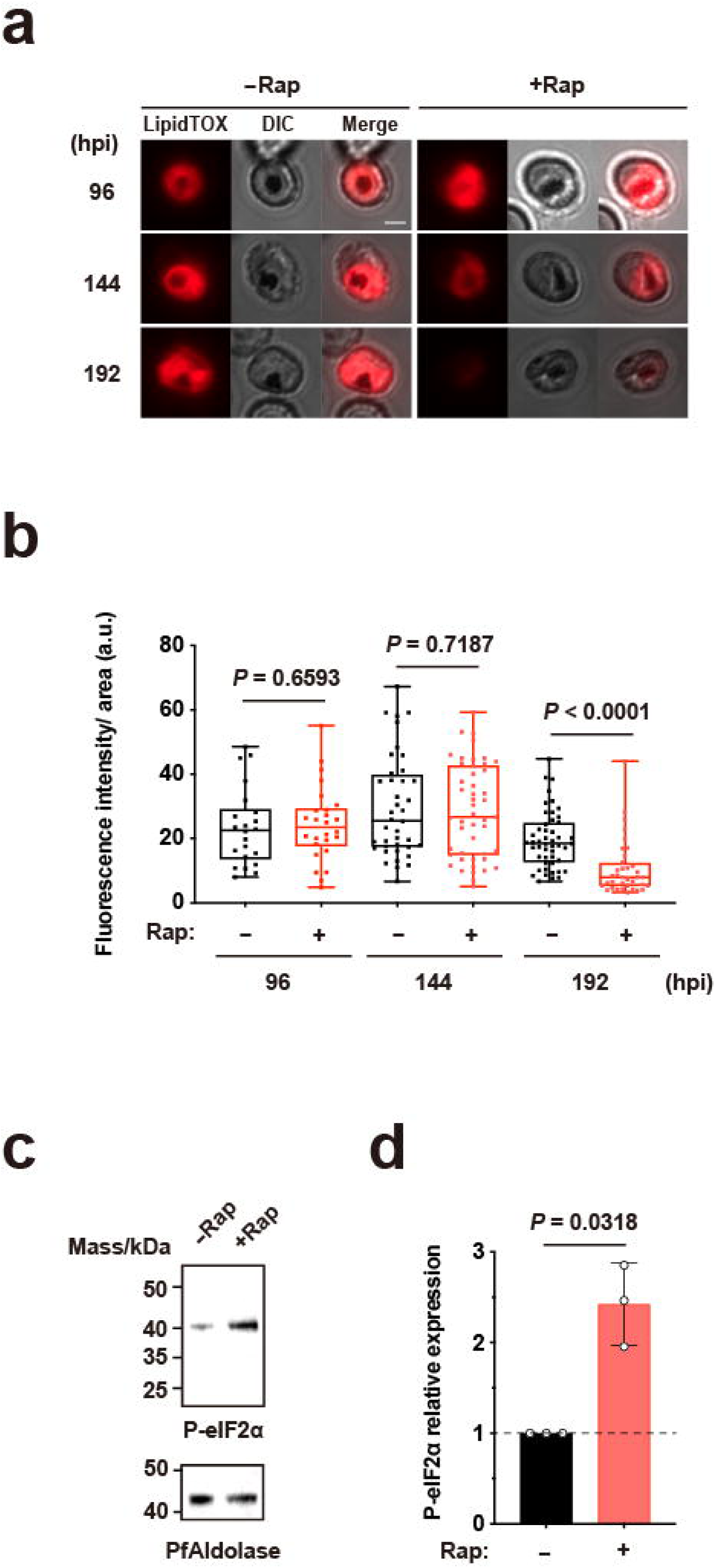
PfDGAT deficiency reduced NL levels and triggered ER stress in parasites. (a) Representative images of LipidTOX-stained +Rap or −Rap *Pfdagt*:LoxPint:HA at the described time points. Scale bars = 2 μm. (b) Boxplots showing the fluorescence intensity of LipidTOX NL staining per area in +Rap or −Rap *Pfdagt*:LoxPint:HA at 96, 144, and 196 hpi. Vertical lines range from the minimum to the maximum values. Horizontal lines represent the 25th percentile (bottom), median (middle), and 75th percentile (top). P values were calculated using Mann–Whitney U test. Experiments were repeated twice. (c) Western blot analysis of phospho-PfeIF2α (P-PfeIF2α) expression in +Rap and −Rap *Pfdagt*:LoxPint:HA after 216 hours of rapamycin treatment. PfAldolase was used as an internal control for the total amount of proteins. Theoretical molecular masses (kDa) of P-PfeIF2α and PfAldolase are 38.0 and 40.0, respectively. Experiments were repeated three times. (d) Bar charts showing the relative expression levels of P-PfeIF2α in +Rap and −Rap *Pfdagt*:LoxPint:HA after 216 hours of rapamycin treatment. Expression levels in the −Rap group were set to 1, and those in the +Rap group were normalized accordingly. P value was calculated using a paired, two-tailed *t*-test. Source data are provided as a Source Data file.

Following the decrease in TAG levels in PfDGAT-deficient parasites, available acyl-CoAs are no longer used for TAG synthesis. This shift results in an abnormal phospholipid-to-NL ratio or a disruption of the equilibrium between free fatty acids (FFA) and acyl-CoAs, ultimately leading to an elevated FFA level in the ER, as reported elsewhere^33^ (Fig. S1, S2). This may trigger ER stress, which could ultimately lead to parasite death, as demonstrated by the growth assay (Fig. 2f). To confirm whether ER stress is induced in *Pfdgat*-deleted parasites, we investigated the phosphorylation levels of *Plasmodium falciparum* eukaryotic initiation factor 2α (PfeIF2α), a marker of ER stress^34^,35. At 216 hpi, +Rap parasites showed higher levels of phosphorylated PfeIF2α (P-PfeIF2α) than −Rap (Fig. 3c), suggesting PfDGAT deficiency promoted ER stress in the parasite.

### PfDGAT deficiency altered the triacylglycerol and phospholipid profiles in the parasite

To investigate the effect of PfDGAT on triacylglycerol and phospholipid profiles, we performed lipidomic analyses on +Rap and −Rap *Pfdgat*:LoxPint:HA parasites at 78 hpi, a time point when the parasites had not yet shown signs of death or morphological abnormalities (Fig. 2g–i). In +Rap parasites, the fractions of 16 TAG species increased relative to the total TAG content (Fig. 4a–c). These elevated TAG species contained at least one polyunsaturated fatty acid (PUFA), suggesting that PUFAs were less susceptible to hydrolysis than saturated fatty acids in the PfDGAT-deficient parasites, although TAG lipase activity has not yet been characterized in terms of its function and substrate preference. In contrast, total TAG levels were comparable between +Rap and −Rap parasites at 78 hpi (Fig. S1a), consistent with the NL staining (Fig. 3a, b).

**Fig. 4.**
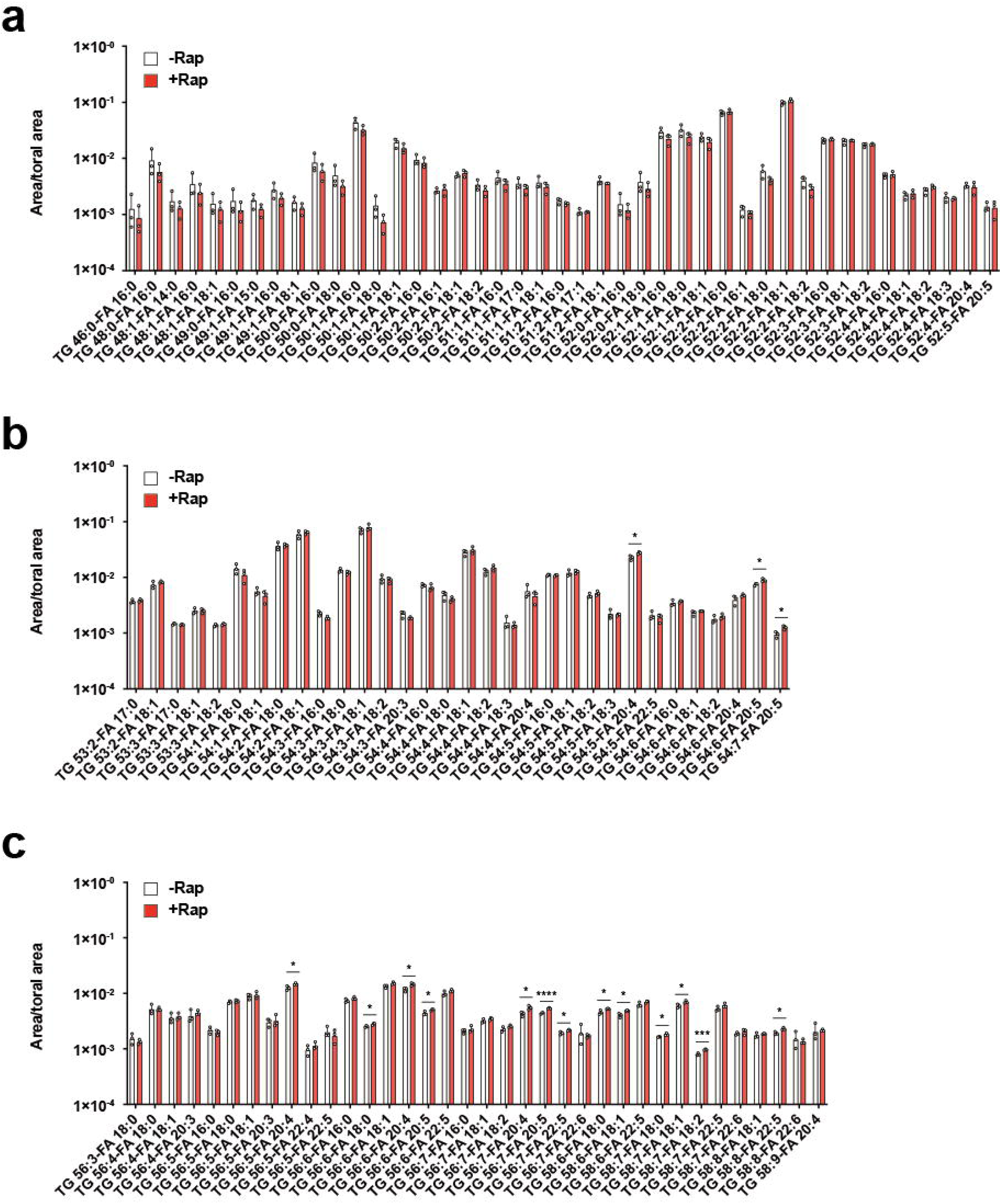
PfDGAT deficiency led to altered TAG profiles in the parasites. (a–c) Bar graphs showing normalized fractions of triacylglycerol (TAG) species against the total area values of TAGs in +Rap and −Rap *Pfdagt*:LoxPint:HA at 78 hpi. On the x-axis, TAGs are denoted as TG. The numbers following an “TG” indicate the total number of carbon atoms and double bounds (e.g., TG 46:0), and the values following “FA” specify the identified acyl chain among the three (e.g., FA 16:0). Data are shown as mean + SD from n = 3 independent biological replicates. P values were calculated using unpaired two-tailed t-test (\**P* < 0.05, \*\**P* < 0.01, \*\*\**P* < 0.005, \*\*\*\**P* < 0.001). Source data are provided as a Source Data file.

Seven classes of phospholipids were detected in our analysis: lysophosphatidylcholine (LPC), phosphatidylcholine (PC), phosphatidylethanolamine (PE), phosphatidylserine (PS), phosphatidylinositol (PI), and sphingomyelin (SM). We quantified the relative abundance of individual molecular species within each phospholipid class in proportion to the total content of that class, thereby identifying which species were affected by *Pfdgat* deletion. In +Rap parasites, most PC species showed a relative increase (Fig. S1a), except for PC(32:0), PC(34:1), and PC(38:9)/PC(e38:1). PE species containing fatty acids with total carbon numbers of 32 to 34 [PE(32–34:n)] were reduced compared to total PE in +Rap parasites, whereas PE (36–49:n) were elevated (Fig. S1b). In contrast, SM(32-36:n) species increased, while SM(40–44:n) decreased in +Rap parasites (Fig. S1e). The relative abundance of PI and PS species was largely unaffected, although the number of detectable species in these classes was lower than in other phospholipid classes.

Following our investigation of relative abundance, we next examined the total abundance of each lipid class, including TAG. The total abundance of all phospholipid classes was markedly elevated in +Rap parasites, whereas TAG levels remained unchanged (Fig. S2a–h). These results indicate that PfDGAT deficiency exerts an impact on phospholipid metabolism before influencing TAG.

### PfDGAT deficiency abolished gametocytogenesis without effecting gametocyte development

Our previous study demonstrated that NL levels were elevated in *P. falciparum* gametocytes^4^. This prompted us to investigate the physiological function(s) of PfDGAT in the gametocyte. To explore the effect of PfDGAT on gametocyte development, gametocytemia was monitored between days 12 and 15 (mid- to late-gametocytogenesis period) following rapamycin treatment on day 9 (Fig. 5a). Although *Pfdgat* was successfully excised from the genome (Fig. 5b), gametocytemia in +Rap parasites remained comparable to that of −Rap parasites (Fig. 5c). Additionally, no obvious differences in gametocyte stage distribution were observed between +Rap and −Rap parasites (Fig. 5d). This suggests that PfDGAT is dispensable for the maturation of gametocytes.

**Fig. 5.**
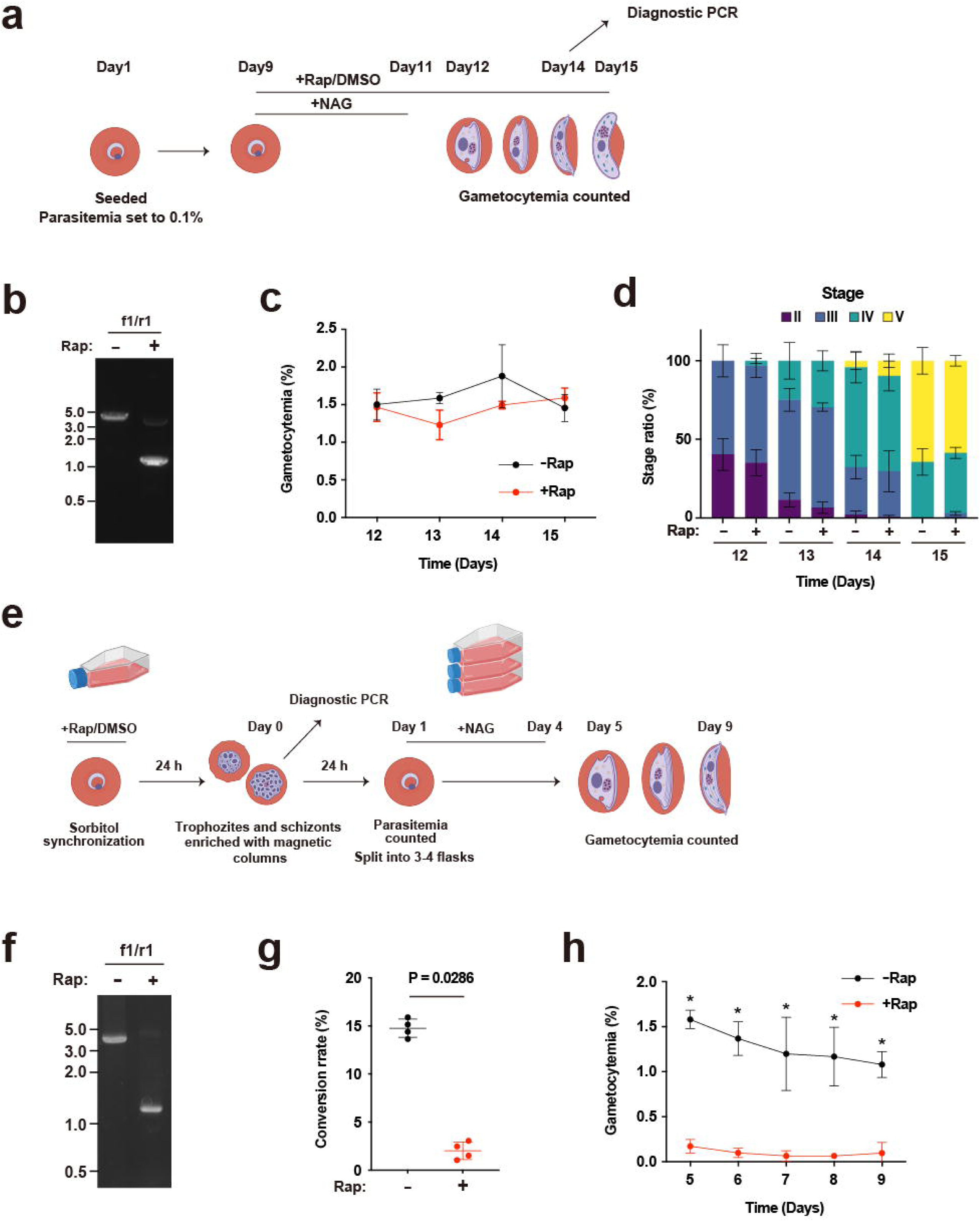
PfDGAT deficiency abolished gametocytogenesis without any effects on gametocyte development. (a) Schematics of experimental procedure to investigate the effect of PfDGAT deficiency on gametocyte development in *Pfdagt*:LoxPint:HA. (b) Conditional knockout of *Pfdagt* in gametocytes was confirmed by PCR on day 14. The size of DNA amplicons shifted from 3704 bp to 1089 bp. (c) Gametocytemia of +Rap or −Rap *Pfdagt*:LoxPint:HA on days 12–15. Data are shown as mean ± SD from n = 3 independent biological replicates. (d) Bar charts showing the gametocyte stage distributions of +Rap or −Rap *Pfdagt*:LoxPint:HA on days 12–15. Data are shown as mean from n = 3 independent biological replicates. (e) Schematics of experimental procedure to investigate the effect of PfDGAT deficiency on sexual conversion rate in *Pfdagt*:LoxPint:HA. (f) Conditional knockout of *Pfdagt* in trophozoites and schizonts was confirmed by PCR on day 0. The size of DNA amplicons shifted from 3704 bp to 1089 bp. (g) Comparison of sexual conversion rates between +Rap and −Rap *Pfdagt*:LoxPint:HA parasites on day 5. P values were calculated using Mann–Whitney U test. Data are shown as mean ± SD from n = 4 independent biological replicates. Experiments were repeated three times with similar results. (h) Comparison of gametocytemia in +Rap and −Rap *Pfdagt*:LoxPint:HA parasites on days 5–9. P values were calculated using two-way ANOVA followed by Šídák’s multiple comparison test. \**P* < 0.0001 (−Rap vs. +Rap *Pfdagt*:LoxPint:HA). Data are shown as mean ± SD from n = 4 independent biological replicates. Experiments were repeated three times with similar results. Fig. 5a and e were created with Biorender.com. Source data are provided as a Source Data file.

Given the elevated levels of intracellular LPC and other phospholipids in +Rap parasites (Fig. S2b), we investigated gametocyte conversion in the PfDGAT-deficient parasite, as extracellular LPC is known to suppress this process^36^. Highly synchronized ring-stage parasites were maintained in the presence or absence of rapamycin for 24 hours (Fig. 5e). Following this treatment, trophozoite- and schizont-stage parasites were collected using magnetic columns, and *Pfdgat* excision was confirmed by diagnostic PCR at day 0 (Fig. 5f). Parasitemia was quantified on day 1, and N-acetylglucosamine (NAG) treatment was initiated and maintained until day 4. From days 5–9, gametocytemia was assessed. We observed that most PfDGAT-deficient parasites failed to develop into gametocytes (Fig. 5g) and that gametocytemia was significantly reduced in +Rap parasites on days 5–9 (Fig. 5h). We quantified the relative expression of AP2-G, a master transcriptional regulator of gametocytogenesis^37,38^, but no differences in transcriptional levels were observed on days 0 and 1 in our quantitative reverse transcription PCR (qRT-PCR) analysis (Fig S3).

### PfDGAT deficiency impaired cytoadhesion of parasites to human brain endothelial cells

Cerebral malaria is primarily attributed to the cytoadherence of parasite-infected erythrocytes to endothelial cells lining the brain microvasculature, a process mediated by the transport of *Plasmodium falciparum* erythrocyte membrane protein 1 (PfEMP1)—a family of adhesive proteins encoded by approximately 60 *var* genes^39^—to the surface of the infected erythrocyte membrane. Given that ER stress has been shown to impair protein trafficking, we investigated the potential role of PfDGAT in parasite adhesion to human brain microvascular endothelial cells (HBMECs). We evaluated the ability of PfDGAT-deficient parasites to adhere to HBMECs under both static and flow conditions (Fig. 6a). +Rap and −Rap parasites were maintained from days 1–6, followed by sorbitol treatment for ring-stage synchronization. Subsequently, one of the two −Rap groups was treated with brefeldin A (BFA), a protein-trafficking inhibitor, to serve as a negative control for cytoadhesion^40^, ^41^. After 18 hours of the treatment, the parasites were transferred either to culture dishes or microfluidic chambers containing cultivated HBMECs. In both conditions, +Rap parasites exhibited significantly reduced cytoadhesion than −Rap, with levels comparable to those of BFA-treated parasites (Fig. 6b, c). This suggests that PfDGAT is involved in parasite cytoadhesion to HBMECs. To elucidate the underlying mechanism responsible for the low cytoadhesive ability of PfDGAT-deficient parasites, protein trafficking was observed in +Rap and −Rap parasites by analyzing the localization of skeleton binding protein 1 (SBP1), a protein exported to Maurer’s clefts^42^, using IFA (Fig. 6d). On days 6 and 8, SBP1 abundance was significantly reduced in PfDGAT-deficient parasites, indicating that PfDGAT plays a potential role in protein trafficking and influences the parasite’s cytoadhesive propensity, possibly through altered PfEXP1 exposure.

**Fig. 6.**
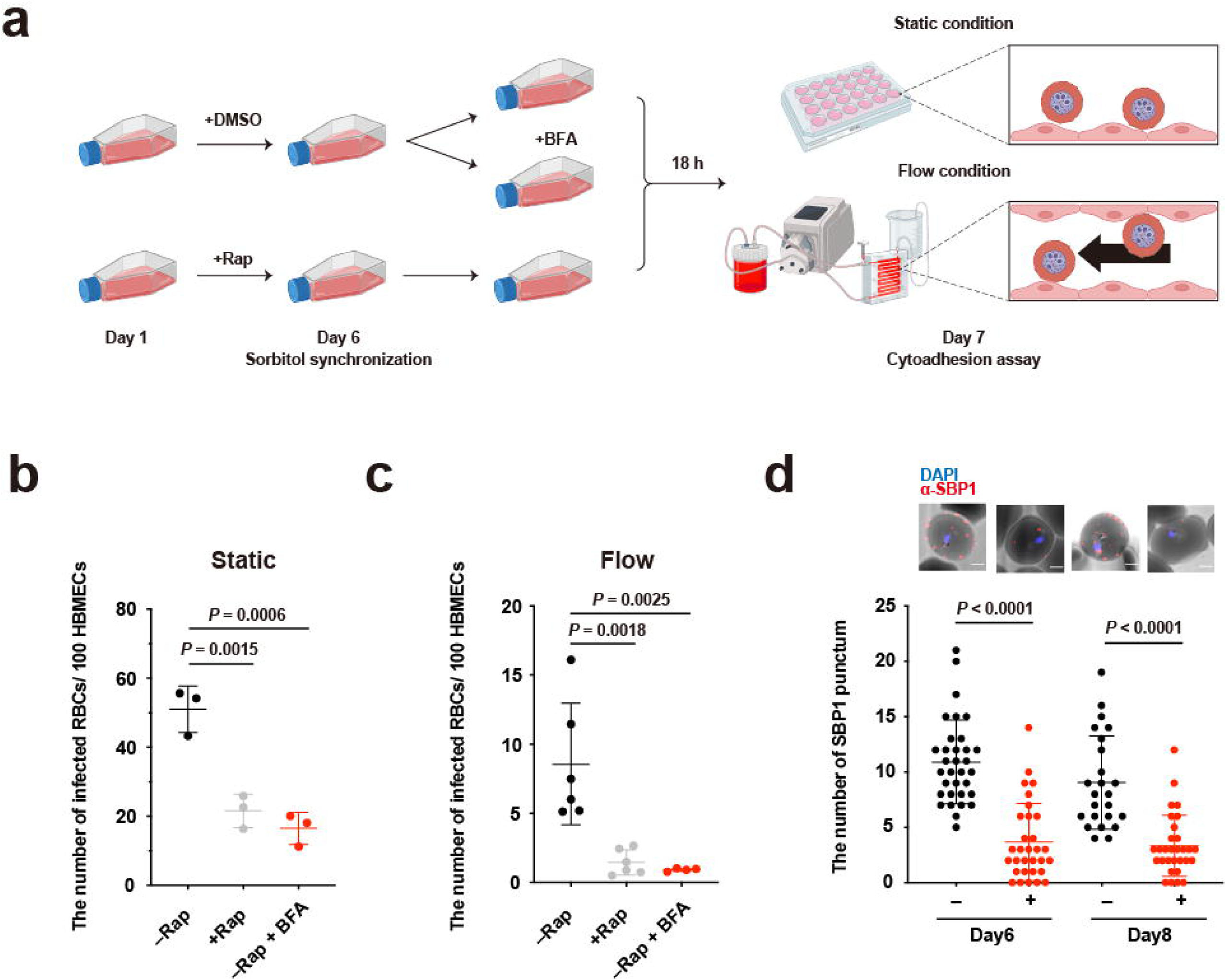
PfDGAT deficiency impaired cytoadhesion of parasites on human brain endothelial cells. (a) Schematics of cytoadhesion assay under static and flow conditions. (b, c) Dot plots showing the number of erythrocytes infected with *Pfdgat*:LoxPint:HA parasites that adhered to 100 human brain microvascular endothelial cells (HBMECs) under static (b) and flow (c) conditions after 6 days of rapamycin treatment. Data are shown as mean ± SD from n = 3 independent biological replicates (b) and from n = 2 independent biological replicates from 3 independent experiments (c). P values were calculated using one-way ANOVA followed by Tukey–Kramer test. BFA, brefeldin. (d) Dot plots showing the number of SBP1 puncta on infected erythrocytes with *Pfdagt*:LoxPint:HA on 6 and 8 days of rapamycin treatment. P values were calculated using one-way ANOVA followed by Tukey–Kramer test. Representative immunofluorescence images are shown above the corresponding plots. Samples were stained for SBP1 with rabbit anti-SBP1 antibody (red) and for nuclei with DAPI (blue). Scale bars = 2 μm. Fig. 6a was created with Biorender.com. Source data are provided as a Source Data file.

**Fig. 7.**
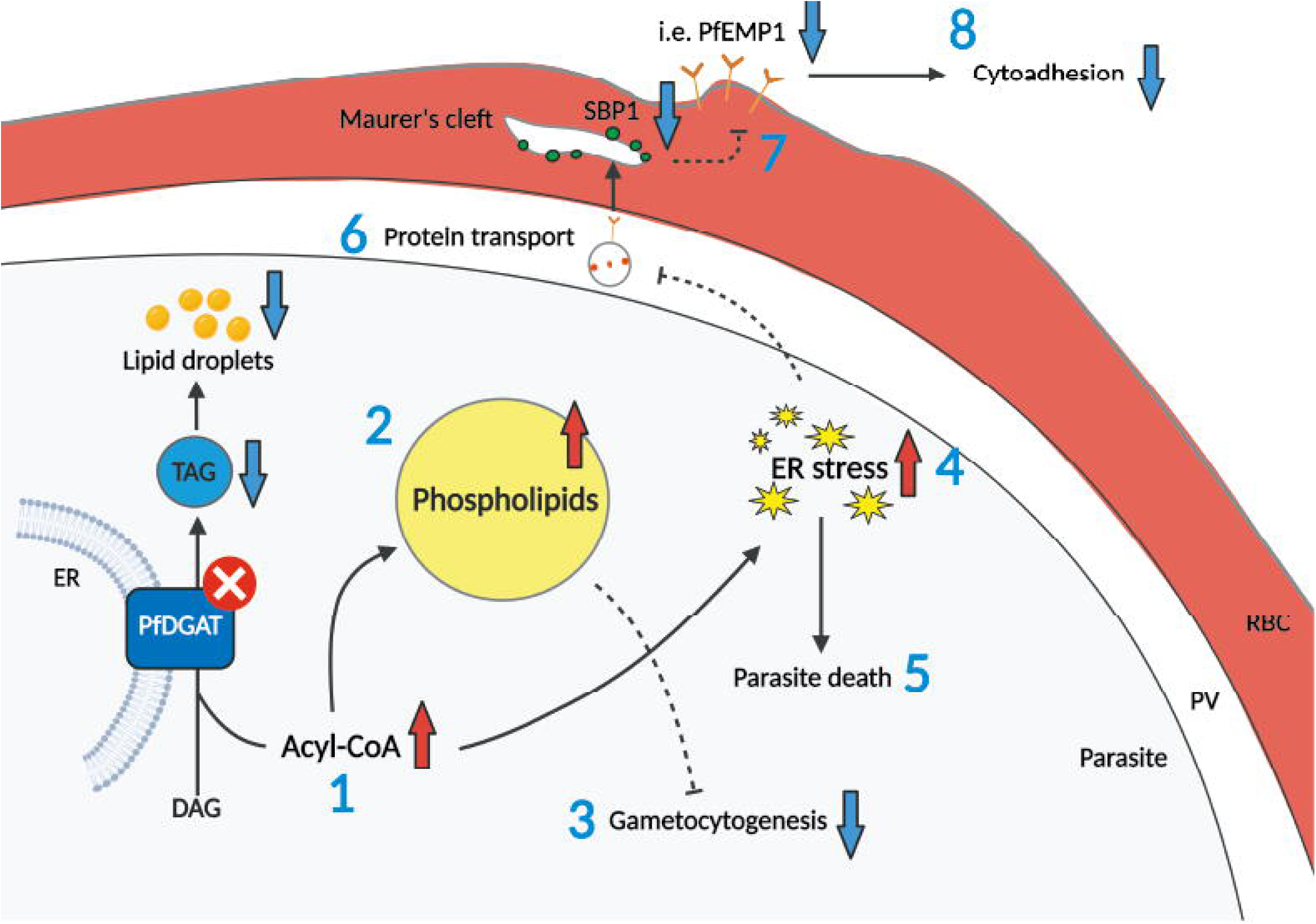
Schematic diagram illustrating the proposed effects of *Pfdgat* deletion on parasites. (1) Deletion of PfDGAT results in elevated free fatty acid (FFA) levels and/or acyl-CoAs due to the decreased incorporation of acyl-CoAs into diacylglycerols (DAGs). (2) Elevated acyl-CoA levels lead to increased phospholipid synthesis. (3) The parasite senses high phospholipid levels as a favorable environment for its asexual replication and therefore suppresses gametocyte differentiation. (4, 5) An excess of FFAs, possibly resulting from increased unused acyl-CoAs, triggers ER stress. A reduction in LDs further exacerbates this stress and ultimately leads to parasite death. (6) ER stress and alterations in membrane lipid composition adversely affect vesicular protein trafficking. (7) Impaired vesicular trafficking reduces the surface presentation of cytoadhesive molecules, such as PfEMP1, thereby diminishing the capacity of parasites to adhere to human endothelial microvascular cells. Created with BioRender.com.

## Discussion

Although PfDGAT has been implicated in LD biogenesis, its biological functions in malaria parasites remain largely uncharacterized. Because the parasite is thought to lack β-oxidation activity^43,44^, which converts the fatty acids released from TAGs into energy sources, it remains unclear how TAGs stored in LDs through PfDGAT activity contribute to the parasite life cycle. In this study, we demonstrated that PfDGAT deficiency results in parasite death, completely blocks sexual differentiation, and severely impairs the cytoadhesive ability of the parasite (Fig. 2e–h, Fig. 5g, h, and Fig. 6b, c), underscoring the pivotal role of this enzyme in the malaria parasite life cycle.

PfDGAT has the potential to become a highly promising drug target in malaria parasites because PfDGAT deficiency adversely affects various aspects of the parasite’s life cycle, as shown in this study. However, a comparative analysis of the amino acid sequences and superimposed structures of PfDGAT and HsDGAT demonstrated that their catalytic motifs and spatial arrangements are similar (Fig. 1a,b). These findings indicate that their substrate preferences and catalytic specificities are likely comparable, suggesting that inhibitors targeting these enzymatic sites would be effective against both enzymes. Indeed, treatment with T863, a selective HsDGAT1 inhibitor targeting the acyl-CoA-binding pocket^45^, resulted in decreased lipid formation and growth arrest in *P. falciparum*^46^, although it remains unclear whether T863 specifically targets PfDGAT. A recent study showed that the N-terminal IDR of HsDGAT1 is essential for dimerization and DGAT activity^25^. While PfDGAT does not possess an discernable IDR at the N-terminus, such a region is predicted to reside between amino acid residues 158 and 220 (Fig. 1a). Although dimerization of PfDGAT was not detected by western blotting under reducing conditions, this IDR may nevertheless play a crucial role in intermolecular interactions and enzymatic activity within the parasite, suggesting that this region could serve as a potential target for PfDGAT-specific drug development.

PfDGAT deficiency probably led to elevated FFAs in the parasite. As excess FFAs cannot be incorporated into DAG in the form of acyl-CoA, this ultimately resulted in reduced TAG levels and fewer LDs (Fig. 3a,b). FFAs are normally either degraded as energy sources via β-oxidation, which is likely inactive in the malaria parasite, or used for phospholipid synthesis as components of the cellular membrane. In PfDGAT-deficient parasites, increased phospholipid levels suggest that the acyl-CoAs not incorporated into DAG are diverted to phospholipid synthesis (Fig. S2). We reasoned that the parasite senses elevated levels of phospholipids, particularly LPC, as a signal of a replication-favorable environment, and thus halts its sexual differentiation, even under gametocytogenesis-inducing conditions (Fig. 5g, h). Excess FFAs have been demonstrated to trigger ER stress^47–49^, as was also observed in PfDGAT-deficient parasites (Fig. 3c). ER stress also elicits adaptive responses, such as the unfolded protein response, which is mediated by the membrane perturbation sensors IRE1 and PERK^50^ and serves to alleviate stress. Because phospholipid levels have been shown to increase under ER stress^51–53^, the increased levels of phospholipid observed in PfDGAT-deficient parasites may reflect not only a diversion of metabolic flux but also a parasite response to ER stress. However, prolonged ER stress ultimately leads to parasite death in PfDGAT-deficient parasites (Fig. 2e–h), as reported previously^54^.

PfDGAT-deficient parasites showed reduced binding to HBMECs in our cytoadherence analysis. Given the decreased levels of SBP1 observed in the Maurer’s clefts of infected erythrocytes, this reduction was likely due to ER stress (Fig. 3c, d), which as previously reported, may impair protein vesicular trafficking in malaria parasites^55,56^. Notably, DGAT1 deficiency has been shown to cause aberrant protein trafficking in human enterocytes^57^, although the underlying mechanism remains still unclear and warrants further investigation. One possible explanation for these observations is that alterations in the balance between TAG and DAG, as well as in the phospholipid composition, under ER stress may contribute to defective protein trafficking. This is plausible because membrane compositions, particularly the levels of PE, PS, and DAG, are known to influence membrane and vesicle fusion processes^9^. In addition, the mislocalization of PfEMP1 may also hinder cytoadherence by disrupting the cellular membrane architecture required for PfEMP1 assembly, possibly due to the excess generation of phospholipids resulting an abnormal phospholipid supply to the membrane (Fig. S1, S2). It is important to note that the cytoadherence assay, including analyses of ER stress and NL quantification, was performed at 96 hpi, a time point when some parasites were already dead, exhibiting morphological abnormalities. To specifically assess the intrinsic effect of PfDGAT deficiency, we selectively counted visually viable parasites with normal morphology under the microscope; however, we cannot rule out the possibility that some dead parasites partially influenced these assay results.

Recently, LDs, generated primarily by DGAT1, have been shown to participate in multiple aspects of cellular metabolism, including the formation of MCSs with other organelles to exchange fatty acids and glycerophospholipids^58–60^. It has been observed that LDs in *P. falciparum* form MCSs with nuclei, digestive vacuoles, the ER, mitochondria, and cytostomes, although their precise functions and associated proteins remain unresolved^46^. Therefore, further investigations of PfDGAT may help reveal unknown aspects of malaria parasite biology.

## Methods

### Ethics statement

The experiments with human erythrocytes were approved by the ethics committee of Nagasaki University (Approved#: 191226226). Human erythrocytes were purchased from the Japan Red Cross Society (Tokyo, Japan) (No: R030027). Informed consent was obtained by the Japan Red Cross Society before the blood samples were purchased (donor information was not disclosed to the purchaser). All ethical regulations relevant to human research participants were followed.

### Parasite culture

*Plasmodium falciparum* NF54::DiCre strain (kindly gifted from Moritz Treeck) was cultured in O^+^ erythrocytes at 5% hematocrit in RPMI-1640 medium (Gibco, Thermo Fisher Scientific, Waltham, MA, USA) supplemented with 5.95 mg/mL HEPES, 0.05 mg/mL hypoxanthine (Sigma-Aldrich, Merck, Darmstadt, Germany), 2 mg/mL sodium bicarbonate (Fujifilm Wako Chemical Corporation, Osaka, Japan), 10 μg/mL gentamicin (Gibco), and 0.5% Albumax II (Gibco) (RPMI-1640 complete media) and maintained at 37 °C in a gas mixture of 5% O_2_, 5% CO_2_, and 90% N_2_, as previously described^4^.

### Generation of *Pfdgat*:LoxPint:HA line

To conditionally knock out *Pfldgat*, we generated a pSLI-*Pfdgat*-LoxP-3×2A-HA plasmid to introduce LoxP sites into *Pfldgat* utilizing an SLI method^31^. To create pSLI-*Pfdgat*-LoxP-3×2A-HA, pSLI-*Pflplat1*-LoxP-3×2A-HA^61^ was linearized at the *Bg*lII and *Xma*I restriction sites. Synthetic DNA containing a LoxP site followed by 3′ recodonized *Pfdagt*, triple influenza hemagglutinin (HA) sequences, a 2A self-cleaving peptide sequence, a neomycin resistance gene cassette, a LoxP site, and a GFP reporter gene was obtained (Azenta Life Sciences, Burlington, MA, USA). The 5’ genomic *Pfdgat* was amplified from the genomic DNA of NF54::DiCre strain with CloneAmp HiFi PCR Premix (Takara Bio, Shiga, Japan) using the primers listed in Table S1. These sequences were introduced into linearized pSLI-*Pflplat1*-LoxP-3×2A-HA using an In-Fusion HD Cloning Kit (Takara Bio).

For plasmid DNA transfection, trophozoite and schizont-stage parasites were collected using MACS magnetic columns (Miltenyi Biotec, Bergisch Gladbach, Germany). Plasmids were electroporated into erythrocytes using a GenePulser Xcell (Bio-Rad, Hercules, CA, USA) with cuvettes (Bio-Rad). Collected parasites were incubated with electroporated erythrocytes in RPMI-1640 complete medium for 48 hours. After 48 hours, parasites were selected with 1 nM WR99210 (Sigma-Aldrich) for 3–4 weeks followed by 400 µg/mL G418 (Fujifilm Wako Pure Chemical Corporation) for 2 weeks. Clonal parasites were obtained by limiting dilution of drug-selected transgenic parasites. To confirm plasmid integration into the genome of clonal parasites, genomic DNA was extracted using a DNeasy Blood & Tissue Kit (QIAGEN, Hilden, Germany) and a diagnostic PCR was performed using the primers listed in Table S1.

### Three-dimensional construction of PfDGAT and HsDGAT1

For the structural overlay of PfDGAT and HsDGAT1, Protein Data Bank files were retrieved from the AlphaFold Protein Structure Database (https://alphafold.ebi.ac.uk/) (PfDGAT; AF-O97295-F1-model_v4, HsDGAT1; AF-O75907-F1-model_v4). The proteins were visualized and superimposed using PyMOL (v3.1.4.1) (Schrödinger, USA).

### Multiple sequence alignment of DGAT1

For multiple alignment of DGAT1 sequences, amino acid sequences were obtained from the NCBI database (https://www.ncbi.nlm.nih.gov/) (*Homo sapiens* DGAT1 [HsDGAT1], NP_036211.2; *Plasmodium falciparum* 3D7 DGAT [PfDGAT], XP_001351293; *Plasmodium vivax* P01 DGAT [PvDGAT], XP_001614490; *Plasmodium knowlesi* strain H DGAT [PkDGAT], XP_002258826; *Plasmodium berghei* ANKA DGAT [PbDGAT], XP_678753; *Toxoplasma gondii* ME49 DGAT [TgDGAT], XP_018638224; *Cryptosporidium parvum* IowaII DGAT [CpDGAT], XP_625364). The alignment was carried out using CLUSTALW (v2.1). Sequence logos were created using WebLogo (v2.8.2) (https://weblogo.berkeley.edu/logo.cgi).

### Growth assay

For growth assays, parasites were synchronized with 5% D-sorbitol (Fujifilm Wako Pure Chemical Corporation) and seeded into 24-well plates. Synchronized parasites were maintained in 1 mL of RPMI-1640 complete medium at 5% hematocrit and treated with either 100 nM rapamycin (Sigma-Aldrich) or an equivalent volume of dimethyl sulfoxide. Parasitemia was monitored daily using a Sysmex XN-30 hematology analyzer (Sysmex Corporation, Hyogo, Japan)^62^ and maintained below 5% by serial passaging. For analysis, membranes of parasitized erythrocytes were permeabilized with CELLPACK DCL (Sysmex Corporation), and parasite nucleic acids were stained with Fluorocell M and Lysercell M (Sysmex Corporation). Growth curves were generated using GraphPad Prism v10.5.0 (GraphPad Software, San Diego, CA, USA).

### Quantification of parasite stage distribution

For the quantification of parasite stage distribution, trophozoite- and schizont-stage parasites were collected using MACS magnetic columns (Miltenyi Biotec). Pelleted samples were resuspended at 2% hematocrit in RPMI-1640 complete medium for 4 hours under continuous agitation at 40 rpm to reduce multiple infections of parasites within a single erythrocyte. After 4 hours of incubation, parasites were tightly synchronized with 5% D-sorbitol (0–4 hours post invasion) and seeded into 6-well plates. Thin blood smears were prepared at each time point and stained with Giemsa′s azur eosin methylene blue solution (Sigma-Aldrich). Parasite staging was performed manually under light microscopy.

### Immunofluorescence assay

Infected erythrocytes were immobilized on poly-D-lysine–coated coverslips (Thorlabs, NJ, USA; poly-D-lysine from MP Biomedicals, CA, USA). After immobilization, samples were fixed with 4% paraformaldehyde (Electron Microscopy Sciences, PA, USA) in phosphate-buffered saline (PBS) for 10 minutes, followed by crosslinking with 50 mM dimethyl suberimidate dihydrochloride (Sigma-Aldrich) in borate buffer for 20 minutes, and quenched with 10 mM glycine in PBS for 30 minutes. Fixed samples were permeabilized with 0.3% Triton X-100 (Fujifilm Wako Pure Chemical Corporation) in PBS and blocked with 1% skim milk in 0.05% Tween 20 (Tokyo Chemical Industry, Tokyo, Japan) in PBS for 60 minutes. Samples were incubated with primary antibodies overnight and with secondary antibodies for 60 minutes, followed by mounting in VECTASHIELD Antifade Mounting Medium with DAPI (Vector Laboratories, CA, USA). Immunostained samples were observed using a 100× objective on an LSM780 confocal laser scanning microscope (Carl Zeiss, Oberkochen, Germany).

### Western blot

For western blot analysis, parasites were collected using MACS magnetic columns (Miltenyi Biotec) and lysed in 0.03% saponin solution for 5 minutes. Collected samples were dissolved in lysis buffer (4% SDS, 0.5% Triton X-100, 25–75 U DNase I [Takara Bio] and cOmplete protease inhibitor cocktail [Roche, Basel, Switzerland] in 0.5% PBS) and denatured with SDS sample buffer containing dithiothreitol. Lysates were boiled at 50 °C for 3 minutes (PfDGAT-HA) or at 95 °C for 10 minutes (Phospho-PfeIF2α), separated on 4%–12% polyacrylamide gels (Invitrogen), and transferred to PVDF membranes. Membranes were blocked with Bullet Blocking One (Nacalai Tesque, Kyoto, Japan), incubated overnight with primary antibodies, then incubated for 30 minutes with horseradish peroxidase (HRP)-conjugated secondary antibodies. Signals were detected using chemiluminescent HRP substrate (Thermo Fisher Scientific) and visualized with WSE-6100 LuminoGraph I (ATTO, Tokyo, Japan).

For internal control detection of PfAldolase, membranes were stripped with WB Stripping Solution Strong (Nacalai Tesque) for 30 minutes, re-blocked with Bullet Blocking One, incubated with HRP-conjugated anti-*Plasmodium* aldolase antibody for 30 minutes, and the signal was developed with chemiluminescent HRP substrate (Thermo Fisher Scientific) for 5 minutes.

### Gametocytogenesis

Gametocyte production was induced by initiating cultures at 6% hematocrit in complete RPMI medium supplemented with 10% human serum (Japan Red Cross, lot #28J0058) and 0.1% parasitemia^4^ (day 1). Hematocrit was reduced to 3% on day 4. Asexual parasites were eliminated by treatment with 50 mM N-acetyl glucosamine (Sigma-Aldrich) to remove asexual parasites on days 9–11 and treated with 100 nM rapamycin (Sigma-Aldrich) from day 10 onward to induce conditional knockout. Gametocytemia and parasite stage distribution were assessed manually from Giemsa-stained blood smears on days 12–15. Media were replaced with pre-warmed and fresh media every day up to day 15. To investigate the gametocyte conversion rate, parasites were synchronized with 5% D-sorbitol and incubated with 100 nM rapamycin for 24 hours. After treatment, trophozoite- and schizont-stage parasites were collected using MACS magnetic columns (Miltenyi Biotec) and maintained with 10–15 times the volume of erythrocytes to parasite for 24 hours (day 0). The parasites were treated with 50 mM NAG (Sigma-Aldrich) on days 1–4. Giemsa-stained blood films were created to count ring-stage parasitemia on day 1 and gametocytemia on days 5–8. The gametocyte conversion rate was calculated by dividing the gametocytemia count on day 5 by the parasitemia count on day 1. For the extraction of genomic DNA from gametocytes, parasites were enriched at the interface between 40% and 60% Percoll layers and subsequently collected using MACS magnetic columns (Miltenyi Biotec). Genomic DNA was extracted using a DNeasy Blood & Tissue Kit (Qiagen), and PCR was performed using the primers listed in Table S1.

### Quantitative reverse transcription PCR

For quantitative reverse transcription PCR (qRT-PCR), infected erythrocytes were suspended in 10 volumes of TRIzol (Invitrogen, Thermo Fisher Scientific, Waltham, MA, USA) relative to erythrocyte volume and stored at −80°C until use. From the suspension, RNA was extracted with a PureLink RNA Mini Kit (Invitrogen), and cDNA was synthesized from 0.1–1 µg of the RNA using a ReverTra Ace qPCR RT kit (Toyobo, Osaka, Japan). Specific DNA regions of AP2-G and the housekeeping gene seryl-tRNA synthetase (PF3D7_0717700) were amplified from the cDNA using a KAPA SYBR Fast qPCR kit (Kapa Biosystems, MA, USA) and a QuantStudio 3 Real-Time PCR System (Applied Biosystems, Thermo Fisher Scientific, Waltham, MA, USA) with the primers listed in Table S1^63^. Relative quantification of the gene expression was performed using the comparative CT method.

### Neutral lipid quantification

For neutral lipid (NL) quantification, parasites were synchronized at 0–4 hpi and maintained in the presence or absence of 100 nM rapamycin until 96, 144, and 192 hpi. NLs were stained with HCS LipidTOX Red (Invitrogen; 1:1000 dilution) for 30 minutes at 37 °C, washed with RPMI-1640 without Phenol Red (Nacalai Tesque), and counterstained with Hoechst 33342 nuclear stain (Sigma-Aldrich) for 5 minutes. Stained samples were observed under a Leica DMi8 inverted microscope (Leica Microsystems, Wetzlar, Germany). Fluorescence signal intensity was analyzed using ImageJ software (National Institutes of Health, MD, USA).

### SBP1 quantification

For SBP1 quantification, the parasite was synchronized with 5% D-sorbitol and maintained in the presence or absence of 100 nM rapamycin. Six and 8 days after the treatments, the infected erythrocytes were immunostained with rabbit anti-SBP1 antibody^42^, followed by Alexa Fluor 594-conjugated goat anti-rabbit antibody. The immunofluorescence signal for SBP1 was visualized using an LSM780 confocal laser scanning microscope (Carl Zeiss) equipped with a Plan-Apochromat 100×/1.40 Oil DIC M27 (numerical aperture = 1.40). The scanned micrographs were processed using ImageJ software (National Institutes of Health). The SBP1 signal was defined using the Mean method on Threshold menu, and subsequently quantified using the Analyze Particles function after applying Dilate to enhance signal continuity and the Watershed algorithm to separate continuous particles.

### Cytoadherence assay

For the static cytoadherence assay, immortalized human brain microvascular endothelial cells (HBMEC) (P10361-IM; Innoprot, Dario, Spain) were cultivated on 13-mm coverslips (Matsunami Glass, Osaka, Japan) in 24-well plates containing Endothelial Cell Medium (P60104; Innoprot, hereafter referred to as HBMEC medium) supplemented with 5% fetal bovine serum (175012; Nichirei Biosciences, Tokyo, Japan). They were maintained at 37 °C in a humidified 5% CO₂ atmosphere and used at 80%–90% confluence on day 7.

HBMEC-panned parasites, obtained as previously reported^64^,65, were maintained at 4% hematocrit with or without 100 nM rapamycin on day 6, followed by their synchronization with D-sorbitol. On day 7, HBMEC-panned parasites were adjusted to 1%–2% parasitemia at 4% hematocrit across groups. Parasite suspension (200 µL per well) was seeded onto HBMEC monolayers in RPMI-1640 complete medium and incubated at 37 °C for 75 minutes. During incubation, the plate was gently agitated every 30 minutes to resuspend the settled red blood cells. After 75 minutes, the medium was removed from each well, and HBMECs were gently washed with RPMI medium. This washing step was repeated five times in total. After the second or third wash, the coverslips were transferred to a new 24-well plate, and the remaining washes were performed. After removing the medium, 2.5% glutaraldehyde (400 μL per well, Sigma-Aldrich) in 1% PBS was added to each well and incubated overnight for fixation. The fixative was then removed, and the samples were stained with Giemsa′s azur eosin methylene blue solution (Sigma-Aldrich) for 15 minutes. The coverslips were covered and air-dried for about 2 days. Finally, they were mounted onto glass slides with the cell-adherent side facing down and secured for microscopic observation.

For the flow cytoadherence assay, HBMECs were seeded onto µ-Slide VI 0.1 microfluidic channel slides (80666; Ibidi, Gräfelfing, Germany). They were maintained at 37 °C in a humidified 5% CO₂ atmosphere and used at 80%–90% confluence on day 7. On day 7, synchronized HBMEC-panned parasites were adjusted to 1%–2% parasitemia at 2% hematocrit across groups. After replacing the HBMEC medium in the channel slides, the parasites were maintained at 38 °C on a hot plate. The parasite suspension was then continuously perfused through the slides at a shear stress of 6 dyn/cm² (flow rate: 6.33 mL/min) for 20 minutes per lane using a peristaltic pump (AC-2110II-3; ATTO), followed by flushing with RPMI-1640 medium for an additional 20 minutes per lane. The suspension was introduced through one inlet connected to silicone tubing (3 mm diameter), while RPMI-1640 medium was simultaneously perfused through the other. After parasite perfusion was completed in each lane, the channel was immediately washed with RPMI-1640 medium. Prior to the introduction of a new sample, RPMI-1640 was perfused through the tubing to wash it. After all lanes had been washed, 2.5% glutaraldehyde in 1% PBS was gently perfused through the slides to achieve fixation, and 20–30 images were acquired per lane.

### Liquid chromatography-mass spectrometry analysis

For liquid chromatography-mass spectrometry analysis, parasites were purified at 78 hpi with a magnetic column, and total lipids were extracted with 50 volumes of 2-propanol relative to the volume of parasite pellets. The extracted lipids were analyzed for phospholipids using a triple quadrupole mass spectrometer coupled to a Nexera liquid chromatography system (LCMS-8050 system, Shimadzu) according to a previously published protocol^66^. Briefly, the extracted lipids were separated on an Acquity UPLC BEH C8 column (1.7 µm, 2.1 x 100 mm, Waters) with a gradient of mobile phase A (5 mM NH_4_HCO_3_ in water), mobile phase B (acetonitrile), and mobile phase C (2-propanol) at a flow rate of 0.35 ml/min. The gradient program (time (A:B:C%)) was as follows: 0 minutes (75:20:5), to 20 minutes (20:75:5), to 40 minutes (20:5:75), to 45 minutes (5:5:90), to 50 minutes (5:5:90), and to 55 minutes (75:20:5). The column oven temperature was set at 47 °C. The eluent was ionized by electrospray ionization, then analyzed in the mass spectrometer in selected reaction monitoring (SRM) mode. The SRM transitions (Q1 and Q3) for the head group of phospholipids were ([M + H]+, 184) for PC, ([M + H]+, neutral loss of 141) for PE, ([M – H]–, neutral loss of 87) for PS, and ([M – H]–, 241) for PI. Peak identification and peak area integration were performed using LabSolutions software (Version 5.99 SP2, Shimadzu) according to the manufacturer’s instructions. The peak area of each phospholipid species was divided by the sum of the total peak area for all detected phospholipids with the same head group and presented as the area ratio.

Triglyceride (TG) levels were measured using previously published LC-MS/MS method with minor modifications^67^. A triple quadrupole mass spectrometer (LCMSlll8060; Shimadzu Co., Kyoto, Japan) coupled to a UHPLC system (Nexera; Shimadzu Co.) was used. The extracted lipids were separated on an Shim-pack Velox C18 column (2.1 × 50 mm, 2.7 μm; Shimadzu Co.) with a gradient of mobile phase A (20 mM NH_4_HCO_2_ in water), mobile phase B (acetonitrile), and mobile phase C (2-propanol) at a flow rate of 0.4 ml/min. The gradient program (time (A:B:C%)) was as follows: 0 minutes (15:15:70), to 0.1 minutes (15:15:70), to 8.4 minutes (5:20:75), to 9 minutes (5:20:75), to 9.1 minutes (15:15:70), and to 11 minutes (15:15:70). The column oven temperature was set at 45°C. The eluent was ionized by electrospray ionization and analyzed in selected reaction monitoring (SRM) mode. For each SRM transition, Q1 was set to the ammonium-adduct parent ion and Q3 to the neutral loss corresponding to a fatty acyl chain. Instrument operation was controlled using Shimadzu LabSolutions LCMS software, version 5.112 (Shimadzu Co.). Species-level TG annotations were expressed as the total number of carbon atoms and the total number of double bonds of the three fatty acyl chains. Observed TG peaks were named as the species-level TG annotation followed by the fatty acyl chain detected as the neutral loss. TG peaks that exhibited a peak height of ≥100,000 in all samples within any group were selected for quantification. Peak areas for peaks with a peak height of ≥30,000 were calculated using Traces software^66^.

### Statistics and data reproducibility

Data normality was checked using the Shapiro–Wilk test. Statistical significance is shown in each graph via a P value. A P value of less than 0.05 was considered statistically significant. Statistical tests were performed using GraphPad Prism (v10.5.0).

## Data availability

All source data and uncropped gel and blotting images are included in a Source Data file.

## Acknowledgments

This work was partially supported by Grants-in-Aid for Scientific Research (KAKENHI), grant numbers 17K08805 (F.T.), 23K06518 (F.T.), 21K15429 (J.F.), and 25K18790 (J.F.); the Japan Institute for Health Security (JIHS) Intramural Research Fund (22T001 to HS); and by a foundation from Shionogi and Co., Ltd. These funders were not involved in the design, experiments, or data analyses in this study. We thank Suzanne Leech, Ph.D., from Edanz (https://jp.edanz.com/ac) for editing a draft of this manuscript.

## Author contributions

Conceptualization: M.Y., J.F., S.M., S.M.T., H.O., and F.T. Data collection: J.F., M.Y., S.M., M.T., S.M.T., and H.O. Data analysis: J.F., M.Y., S.M., M.T., S.M.T., H.O., and F.T. Funding acquisition: J.F., and F.T. Supervision: J.F., and F.T. Writing – original draft: M.Y., J.F., and F.T. Writing – review & editing: M.Y., J.F., S.M., S.M.T., M.T., H.O., T.S., Y.O., K.Y., E.T., D.K.I., K.K., and F.T. Resources: F.T., K.Y., and K.K.

