## Supplementary material for "*Plasmodium falciparum* diacylglycerol acyltransferase maintains phospholipid homeostasis to regulate sexual differentiation, ER stress, and cytoadhesion": Fig. S, Table S

**Table S1. Primers used in this study**

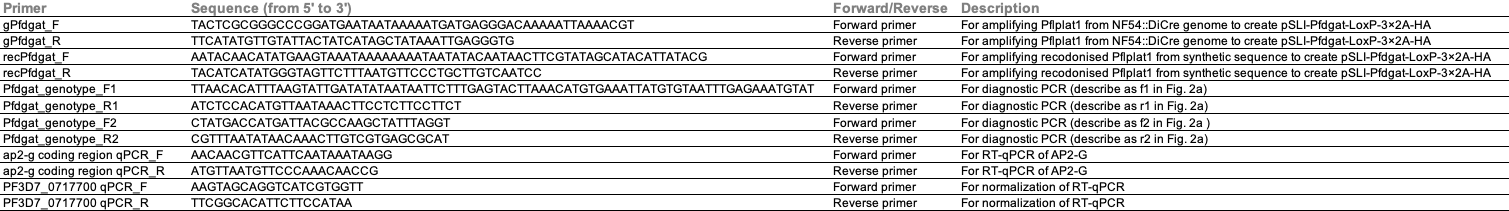

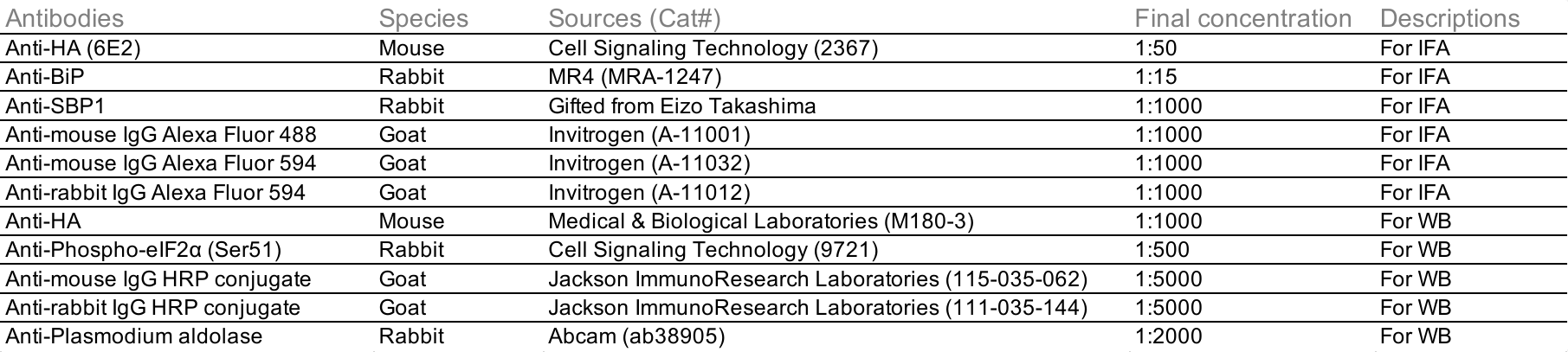
**Table S2. Antibodies used in this study**

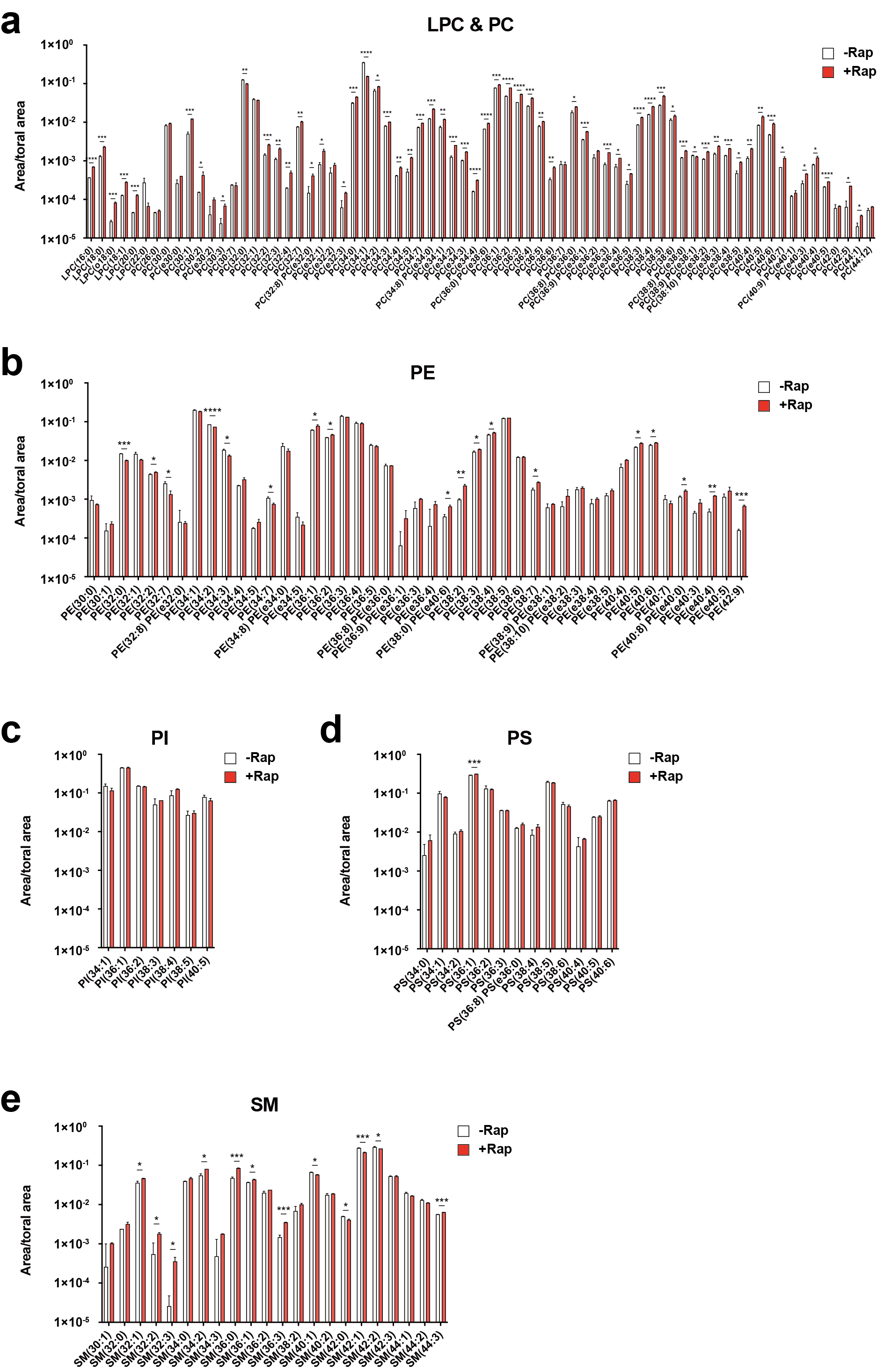

**Fig. S1. *Pfdgat* deficiency affected phospholipid profiles in parasites**

(a–e) Bar graphs showing normalized fractions of lysophosphatidylcholine and phosphatidylcholine (LPC & PC) (a), phosphatidylethanolamine (PE) (b), phosphatidylserine (PS) (c), phosphatidylinositol (PI) (d), and sphingomyelin (SM) (e) species against the total area values within the same classes of phospholipids in +Rap and −Rap *Pfdagt*:LoxPint:HA at 78 hpi. Data are shown as mean + SD from n = 2 independent biological replicates. P-values were calculated using unpaired two-tailed t-test (**P* < 0.05; ***P* < 0.01, ****P* < 0.005, *****P* < 0.001). Source data are provided as a Source Data file.

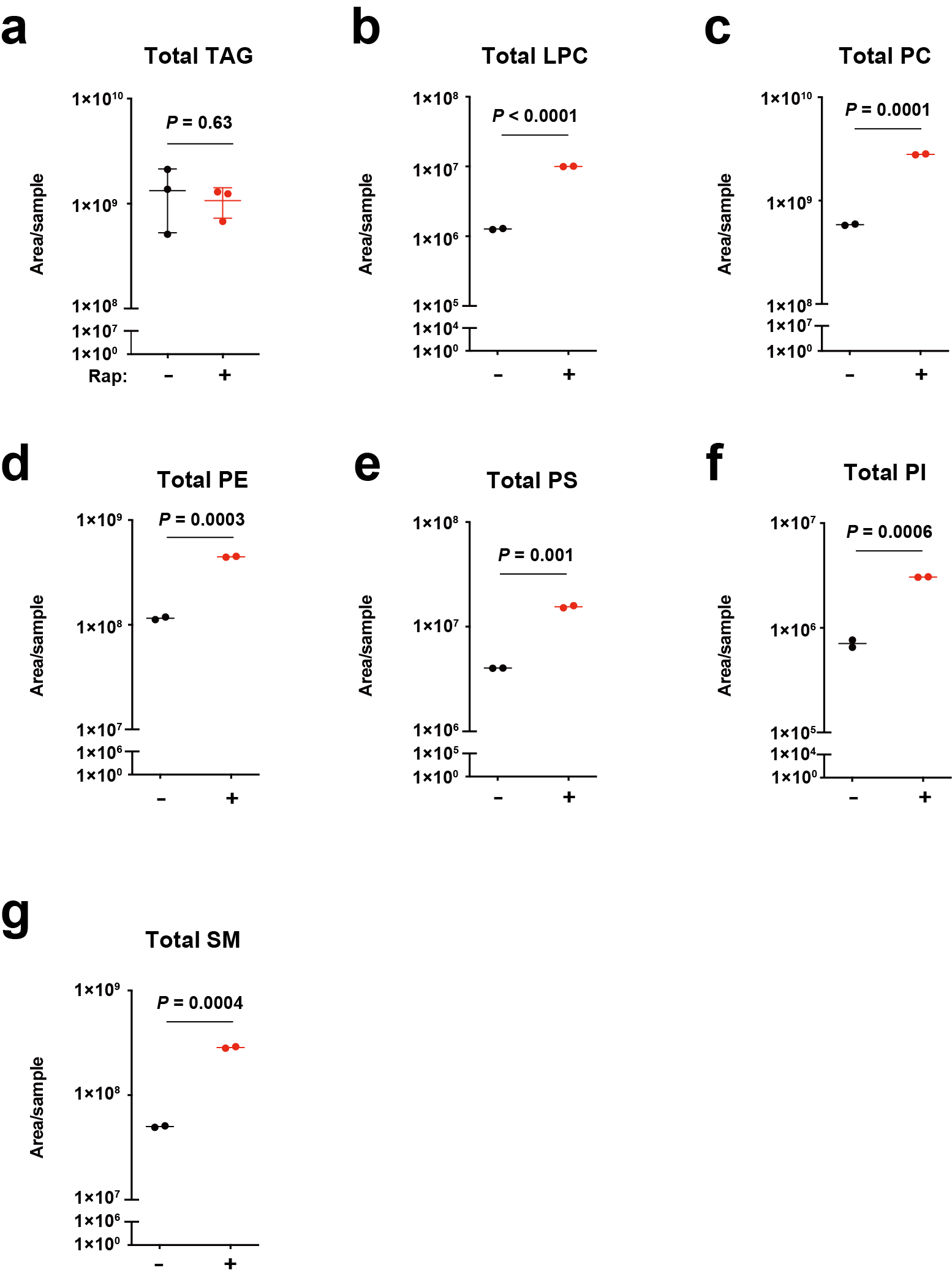

**Figure S2.**

(a–e) Dot plots showing the total area values of TAG (a), LPC (b), PC (c), PE (d), PS (e), PI (f), and SM (g) in −Rap and +Rap *Pfdagt*:LoxPint:HA at 78 hpi. Extracted lipids from an equal number of parasites were analyzed by LC-MS/MS. Data are shown as the mean ± SD from n = 3 independent biological replicates (a) and the mean from n = 2 independent biological replicates (b–h). P-values were calculated using unpaired two-tailed t-test (**P* < 0.05; ***P* < 0.01, ****P* < 0.005, *****P* < 0.001). Source data are provided as a Source Data file.

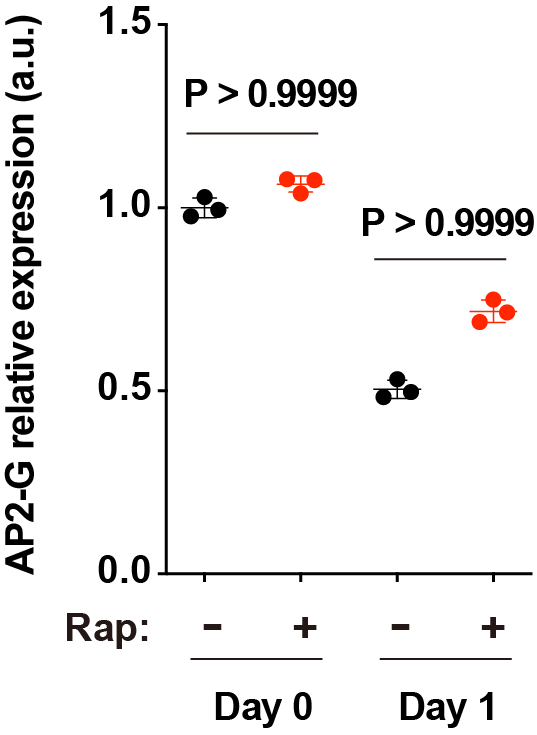

**Figure S3.**

Comparison of relative AP2-G expression between −Rap and +Rap *Pfdagt*:LoxPint:HA on days 0 and 1. The expression level in the −Rap parasite on day 0 was normalized to 1.0. P-values were calculated using Kruskal–Wallis test followed by Dunn's test. Data are shown as mean ± SD from n = 3 independent biological replicates. Experiments were repeated twice with similar results. Source data are provided as a Source Data file.
